# Dual targeting of hepatocyte DGAT2 and stellate cell FASN alleviates nonalcoholic steatohepatitis in mice

**DOI:** 10.1101/2023.07.05.547848

**Authors:** Batuhan Yenilmez, Shauna Harney, Chloe DiMarzio, Mark Kelly, Kyounghee Min, Dimas Echeverria, Brianna M. Bramato, Samuel O. Jackson, Keith Reddig, Jason K. Kim, Anastasia Khvorova, Michael P. Czech

## Abstract

Nonalcoholic steatohepatitis (NASH) is a malady of multiple cell types associated with hepatocyte triglyceride (TG) accumulation, macrophage inflammation, and stellate cell-induced fibrosis, with no approved therapeutics yet available. Here, we report that stellate cell fatty acid synthase (FASN) in de novo lipogenesis drives the autophagic flux that is required for stellate cell activation and fibrotic collagen production. Further, we employ a dual targeting approach to NASH that selectively depletes collagen through selective stellate cell knockout of FASN (using AAV9-LRAT Cre in FASN^fl/fl^ mice), while lowering hepatocyte triglyceride by depleting DGAT2 with a GalNac-conjugated, fully chemically modified siRNA. DGAT2 silencing in hepatocytes alone or in combination with stellate cell FASNKO reduced liver TG accumulation in a choline-deficient NASH mouse model, while FASNKO in hepatocytes alone (using AAV8-TBG Cre in FASN^fl/fl^ mice) did not. Neither hepatocyte DGAT2 silencing alone nor FASNKO in stellate cells alone decreased fibrosis (total collagen), while loss of both DGAT2 plus FASN caused a highly significant attenuation of NASH. These data establish proof of concept that dual targeting of DGAT2 plus FASN alleviates NASH progression in mice far greater than targeting either gene product alone.

## Introduction

Nonalcoholic fatty liver disease (NAFLD) is a malady associated with excess triglyceride accumulation within hepatocytes (hepatic steatosis) and affects nearly 1 billion people globally ^1^. Recent data suggest that by 2040, more than half the adult population will be afflicted by NAFLD, a 43.2% increase from the 2020 prevalence ^2, 3^. Presented by itself, NAFLD can be relatively benign; however, it is often associated with comorbidities such as obesity and type 2 diabetes and therefore varies in severity and impact on health. Importantly, it is estimated that 20-25% of NAFLD patients will progress to develop nonalcoholic steatohepatitis (NASH), characterized by hepatic steatosis in conjunction with severe inflammation and fibrosis ^4–9^. NASH is largely driven by proinflammatory immune cells such as macrophages and Kupffer cells and by stellate cells that secrete collagens and other extracellular matrix proteins ^5, 7, 10, 11^. While lifestyle improvements such as weight loss can alleviate NASH, behavioral modifications are often challenging to execute and have not yielded a definitive solution. Furthermore, clinical trials of dozens of potential therapeutics for NASH have failed and no current treatment is approved for this disease.^2,6,10,12–15^

Evidence has emerged supporting the hypothesis that hepatic steatosis is the major independent initiator of NASH. Multiple human genome-wide association studies show strong links between the incidence of NASH and single-nucleotide polymorphisms in genes specifically related to lipid metabolism, such as *PNLP3, TM6SF2, LYPLAL1, GCKR*, and *PPP1R3.* Not only do these gene variants track with steatosis, but also with steatohepatitis and hepatic fibrosis ^15–23^. Based on these findings, our laboratory ^24^ and others ^25–30^ have investigated the potential of diacylglycerol O-acyltransferase 2 (Dgat2), a crucial enzyme in the synthesis of triglycerides ^31, 32^, as a therapeutic target. Some alleviation of NASH by inhibiting liver DGAT2 has been achieved in mice ^25, 27, 29, 30^ and humans ^25, 28^. In our studies, subcutaneous injection of a fully chemically modified, small interfering RNA (siRNA) targeting Dgat2 (denoted Dgat2-1473) significantly reduced the fatty liver phenotype in a NASH mouse model with obesity, as detected by reduced triglyceride accumulation (>85%, p < 0.0001). However, this improvement in steatosis did not translate into a similar reduction of liver fibrosis, as collagen proteins and expression of genes related to inflammation remained elevated in sdDgat2-injected NASH mice ^24^. Thus, these and other data ^26^ suggest that inhibiting triglyceride synthesis alone is not sufficient to fully reverse NASH, but rather targeting one or more genes in the pathways of inflammation and fibrosis may also be necessary ^8, 33, 34^.

One promising set of candidate genes that control stellate cell fibrosis and could be targeted in combination with hepatocyte DGAT2 include acetyl-CoA carboxylase (ACC), and fatty acid synthase (FASN) in the de novo lipogenesis pathway (DNL). Selective inhibitors of ACC and FASN were found to significantly decrease both procollagen synthesis and DNL in cultured LX2 stellate cells ^35^, suggesting a surprising relationship between the lipogenesis and fibrosis pathways in these cells. In rat models of NASH, inhibiting both DGAT2 and ACC enzymes reduced liver fat and collagen levels as well as circulating ALT/AST enzymes, although some elevation of serum triglyceride (TG) was observed upon ACC inhibition ^26^. In addition, major inflammatory pathways were attenuated by FASN blockade through diminished production of Th17 cells in human primary peripheral blood mononuclear cells (PBMCs) *in vitro* ^36^. Results in preclinical models ^36^ and two recent clinical trials ^37, 38^ using FASN inhibitors have achieved significant decreases in NASH indicators without detectable complications or increased plasma TGs. Although early-stage and preliminary, these advances indicate that targeting de novo lipogenesis may have significant, direct effects on the fibrosis aspect of NASH as well as lipid accumulation. However, the mechanism through which DNL regulates stellate cell activation has remained a key unresolved question in the field.

Based on the above considerations, the present studies had two aims: First, we addressed the mechanism by which FASN is required for stellate cell activation. A major clue came from our recent report ^39^ indicating that DNL and FASN fuels autophagy flux in cells by providing fatty acids in phospholipids for rapid expansion of autophagic membranes. Autophagy is known to be required for stellate cell activation ^40–43^. We show here that blockade of palmitate synthesis by FASN inhibition indeed blunts the activation of stellate cells and their synthesis of collagen in association with attenuation of autophagy progression. A second aim was to test whether selectively inhibiting hepatocyte DGAT2 and stellate cell FASN maximally alleviates NASH development in mice. Here we report proof of concept that such dual targeting is indeed a powerful strategy for alleviating NASH, using a choline-deficient high-fat diet (CDAHFD) mouse model. Overall, our data demonstrate that DGAT2-FASN dual-targeting in the liver can alleviate both steatosis and fibrosis in a well-validated mouse model of NASH.

## Results & Discussion

We first tested the hypothesis that FASN promotes Tgf-β-driven activation of human stellate cells *in vitro*. To investigate the role of palmitate production in stellate cell activation, LX2 human stellate cells were treated with a validated, potent, and specific small molecule inhibitor of FASN, TVB-3664 ^36, 44, 45^, at a final concentration of either 1nM or 100nM for 4 hours. Cells were then subjected to 5ng/ml Tgf-β, a strong stimulator of stellate cell activation ^46^. Forty-eight hours later, the morphology of activated stellate cells, with increased cell size and elongation of spindle-like structures ^46^, was easily observed in Tgf-β treated cells. In contrast, TVB-3664 severely inhibited this morphological change in LX-2 cells that were exposed to Tgf-β (Fig.1A). Moreover, FASN inhibition also blocked Tgf-β-induced type 1 collagen protein expression, aligning with a previous publication^35^ (Fig.1B). Surprisingly, this reduction of collagen 1 protein in response to FASN inhibition correlates with reduced phosphorylated SMAD3 protein, a major driver of collagen production. Taken together, these results suggest that blocking palmitate synthesis attenuates downstream elements in Tgf-β signaling that promote fibrosis, as supported by the reduction in mRNA levels of Col1a1 (Fig 1B) and Acta2(Fig 1C) by TVB-3664.

**Figure 1:**
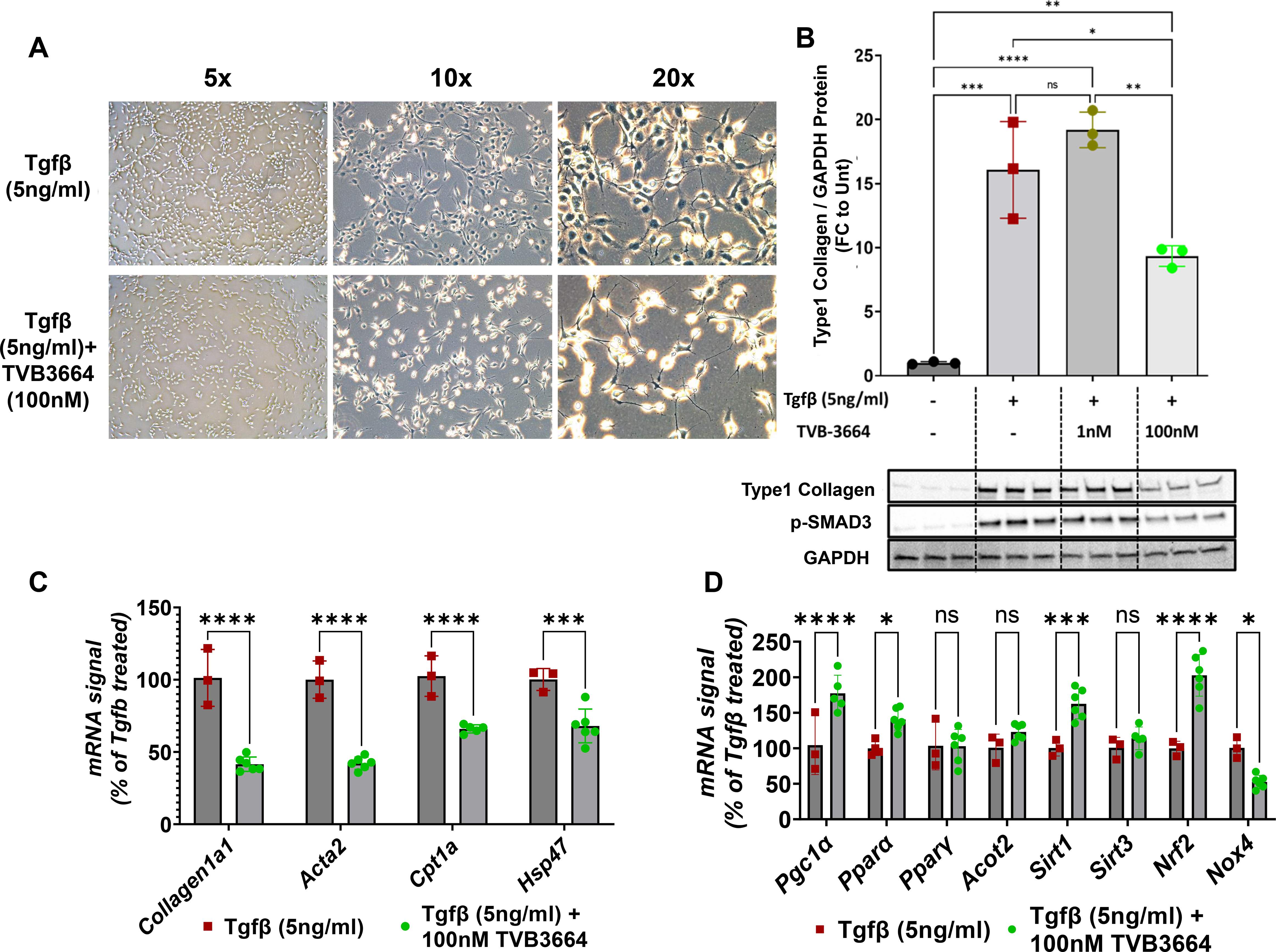
FASN inhibition blunts human stellate cell activation and collagen production. LX2 human stellate cell line was plated in 6 well plates and serum-starved overnight. After 4-hour treatment with 1nM or 100 nM FASN inhibitor (TVB-3664), stellate cell activation was initiated with the addition of recombinant human Tgf-β (5ng/ml final concentration). (A) Cell morphology difference upon FASN inhibition (B) Protein levels of collagen type-1 and phosphorylated Smad3 (C) Fibrotic gene expression changes (D) Oxidative stress and mitochondrial function related gene expression changes (ns: Not significant, *: p<0.05, **:p<0.005, ***:p<0.0005, ****:p<0.00005)

Further analysis of these experiments showed that FASN inhibition leads to a reduction in expression of several genes that have been associated with NASH, including Cpt1a ^47^ and Hsp47^48^, consistent with a more quiescent stellate cell phenotype (Fig.1C). Strikingly, FASN inhibition in Tgf-β activated stellate cells induced remarkable upregulation of Pgc-1α, Pparα, Sirt1, and Nrf2 (Fig 1D), suggesting possible overall improvements in mitochondrial biogenesis as well as a reduction in oxidative stress. Interestingly, Nrf2 has previously been reported to have an inhibitory role in profibrotic SMAD2/3 phosphorylation in stellate cells ^49^. Our findings related to p-SMAD3 levels (Fig.1B) and increased Nrf2 gene expression upon FASN inhibition are consistent with a strong anti-fibrotic effect of TVB-3664. Upregulation of Nrf2 was also correlated with the reduction in expression of Nox4, a major enzyme in the ROS production pathway ^50^, supporting a possible lower oxidative stress state upon FASN inhibition in activated stellate cells (Fig.1D).

Although it is established that inhibition of DNL enzymes blunts stellate cell activation, the underlying mechanisms were unknown. To investigate how inhibition of DNL blocks stellate cell activation, we developed fully chemically modified siRNAs, targeting the human *FASN* transcript. These compounds were designed and synthesized as previously described ^24, 51–53^ (Fig.2A). To identify *FASN* sequences susceptible to potent silencing capability *in vitro*, several test sequences targeting different regions of human *FASN* mRNA were generated by a custom algorithm designed to optimize predicted silencing efficiency of chemically modified RNAs. This panel of compounds was synthesized and initially screened for silencing efficacy in LX2 human stellate cells by direct addition to the culture medium to a final concentration of 1.5 µM. Silencing effects on levels of *FASN* mRNA and housekeeping (*B2M*) gene mRNA after 48 h of treatment was assessed with the QuantiGene assay. Six compounds achieved greater than 70% silencing in the initial screen (Fig.2B). To investigate the potency of sequences identified in the initial screening, dose-response relationship assays were carried out with LX2 cells treated with 8 concentrations of either compound 2314, 5428,8323 or 8324 (1.5 µM to 0.023 µM) for 48h. These potency tests revealed IC50 values of 111.2 nM, 49.1 nM, 147 nM, and 20.4 nM compounds 2314, 5428,8323, or 8324 for silencing human *FASN* mRNA, respectively (Fig.2C).

**Figure 2:**
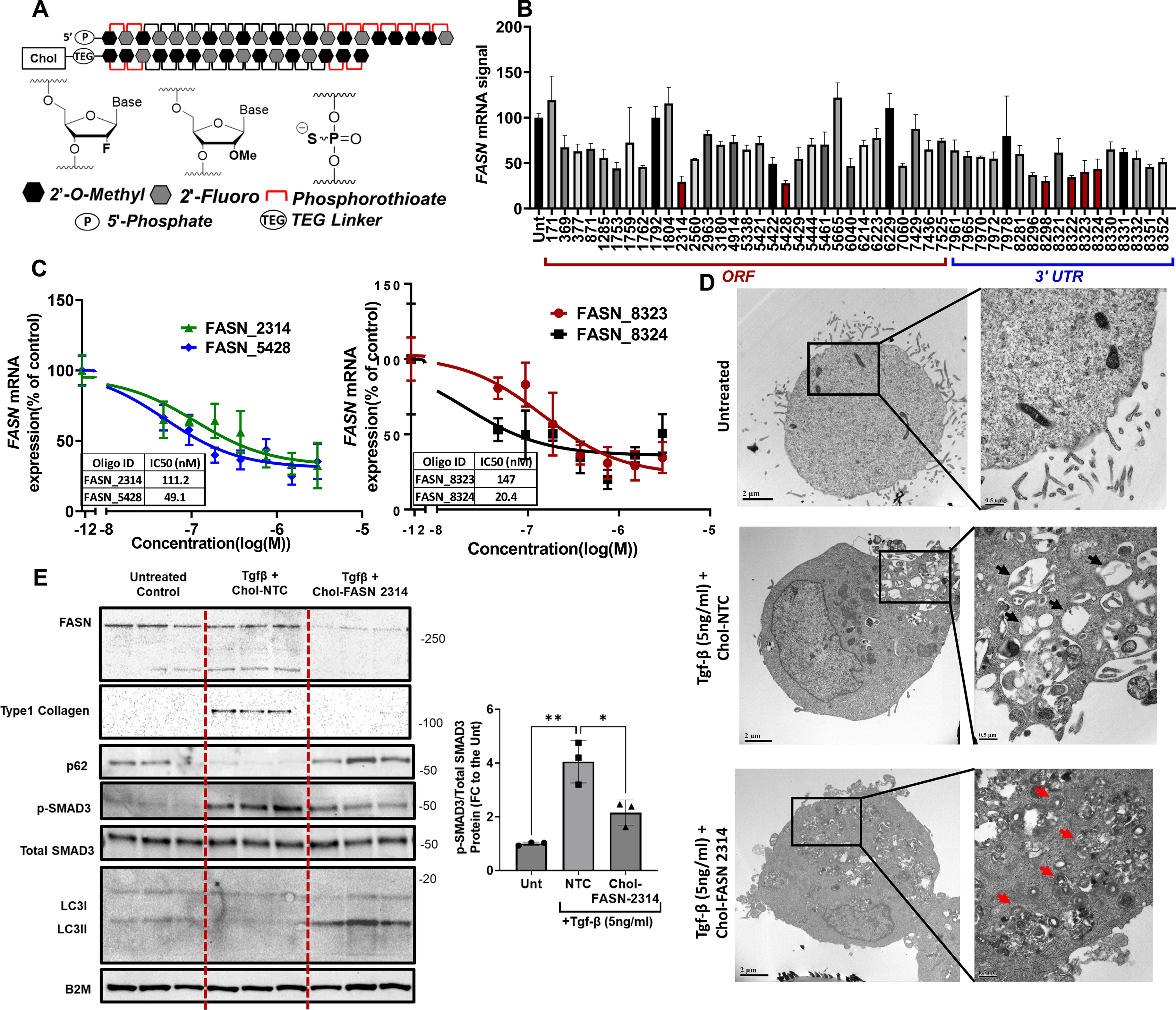
Human *FASN* silencing blunts the activation of cultured human LX2 cells in association with inhibition of autophagic flux. **(A)** Representative cartoon of the chemically modified, cholesterol-conjugated siRNAs that were used for *in vitro* screenings. **(B)** LX2 human stellate cells were treated with siRNA compounds (1.5 µM) for 48 hrs prior to analysis of *FASN* mRNA levels to identify the most potent siRNA sequences **(C)** Dose-response relationships for the lead siRNAs that showed the strongest silencing. The IC50 values were determined by using 8-point serially diluted concentrations of the compounds starting from 1.5 µM. **(D)** TEM images, highlighting autolysosomes (black arrow) and autophagosomes (red arrows) of stellate cells which are Untreated, Tgf-β plus NTC treated, and Tgf-β plus cholesterol conjugated FASN-2314 treated **(E)** Immunoblotting analysis of fibrotic and autophagic pathway markers in stellate cells in a representative experiment with 3 replicates. (ns: Not significant, *: p<0.05, **:p<0.005, ***:p<0.0005, ****:p<0.00005)

Additionally, transmission electron microscopy imaging was utilized to view cellular changes in response to *FASN* silencing (Fig 2D). LX2 human stellate cells were plated in 6 well plates and serum-starved overnight. After 6-hour pre-treatment with either Cholesterol conjugated non-targeting control compound (Chol-NTC) or Cholesterol conjugated FASN-2314 (Chol-FASN-2314), stellate cell activation was initiated with the addition of recombinant human Tgf-β (5ng/ml final concentration). Forty-eight hours later, cells were fixed and sectioned for transmission electron microscopy (TEM). The TEM imaging shown in Fig 2D is in line with previous findings showing increased numbers of autolysosomes during stellate cell activation ^40–43^, indicated with black arrows, in Tgf-β + Chol-NTC treated cells. Strikingly, TEM imaging also revealed that FASN silencing in stellate cells resulted in the accumulation of autophagosomes, indicated with black arrows (Fig.2D), similar to our previous work in adipocytes ^39^. To investigate this issue further, we assessed autophagic activity markers, LC3, and p62, along with fibrogenic markers under the same conditions (Untreated, Tgf-β + Chol-NTC and Tgf-β + Chol-FASN-2314). A strong FASN protein silencing was achieved with Chol-FASN-2314 and this led to strong inhibition of type 1 collagen protein expression and p-Smad3 protein levels (Fig.2E), similar to pharmacological inhibition of FASN shown in Fig.1. Also, p62 and LC3II protein levels were decreased upon activation of stellate cells in Tgf-β + Chol-NTC group compared to non-activated stellate cells (Fig.2E), suggesting increased autophagosome degradation and flux^54^. This conclusion is also supported by the increased number autolysosome presence in the Tgf-β + Chol-NTC group in Fig.2D (Black arrows). *FASN* silencing resulted in a marked increase in levels of LC3II and p62 (Fig.2E), suggesting the autophagosome-lysosome fusion is blocked under these conditions, which is also supported by increased accumulation of autophagosomes in Fig.2D (Red arrows). Altogether, these data strongly support the concept that DNL inhibition blocks fibrogenesis in stellate cells through blockade of autophagic flux that is known to be required for stellate cell activation.

We next investigated the effects of single and dual hepatic targeting of FASN and DGAT2 in mice placed on a NASH-inducing diet (Fig 3). The aim of these studies was two-fold: 1.) to test the effects of selective depletion of FASN in hepatocytes vs liver stellate cells *in vivo*, and 2.) to test the effects of these selective FASNKO perturbations in combination with DGAT2 depletion in hepatocytes *in vivo*. In order to induce such cell-selective FASNKO and DGAT2-depleted mice, FASN^fl/fl^ mice were injected with the following reagents: non-targeting control sdRNA plus control AAV (denoted NTC mice); hepatocyte selective sdDgat2-1473 (sdDgat2) compound for DGAT2 silencing ^24^; AAV8-TBG-Cre for hepatocyte selective FASN KO (FASN HepKO); AAV9-LRAT-Cre for stellate cell selective FASNKO (FASN StelKO); both AAV8-TBG-Cre and AAV9-LRAT-Cre for hepatocyte plus stellate cell-specific FASNKO (FASN HepKO + StelKO). Combinations of these injections were also used to deplete both FASN and DGAT2 in selective cell types, as depicted in Figure 3A. The sdDgat2 compound was injected once subcutaneously, while AAVs were injected once intravenously. One week following injections, mice were placed on a choline-deficient high-fat diet (CDHFD) for 7 weeks (Fig.3A).

**Figure 3:**
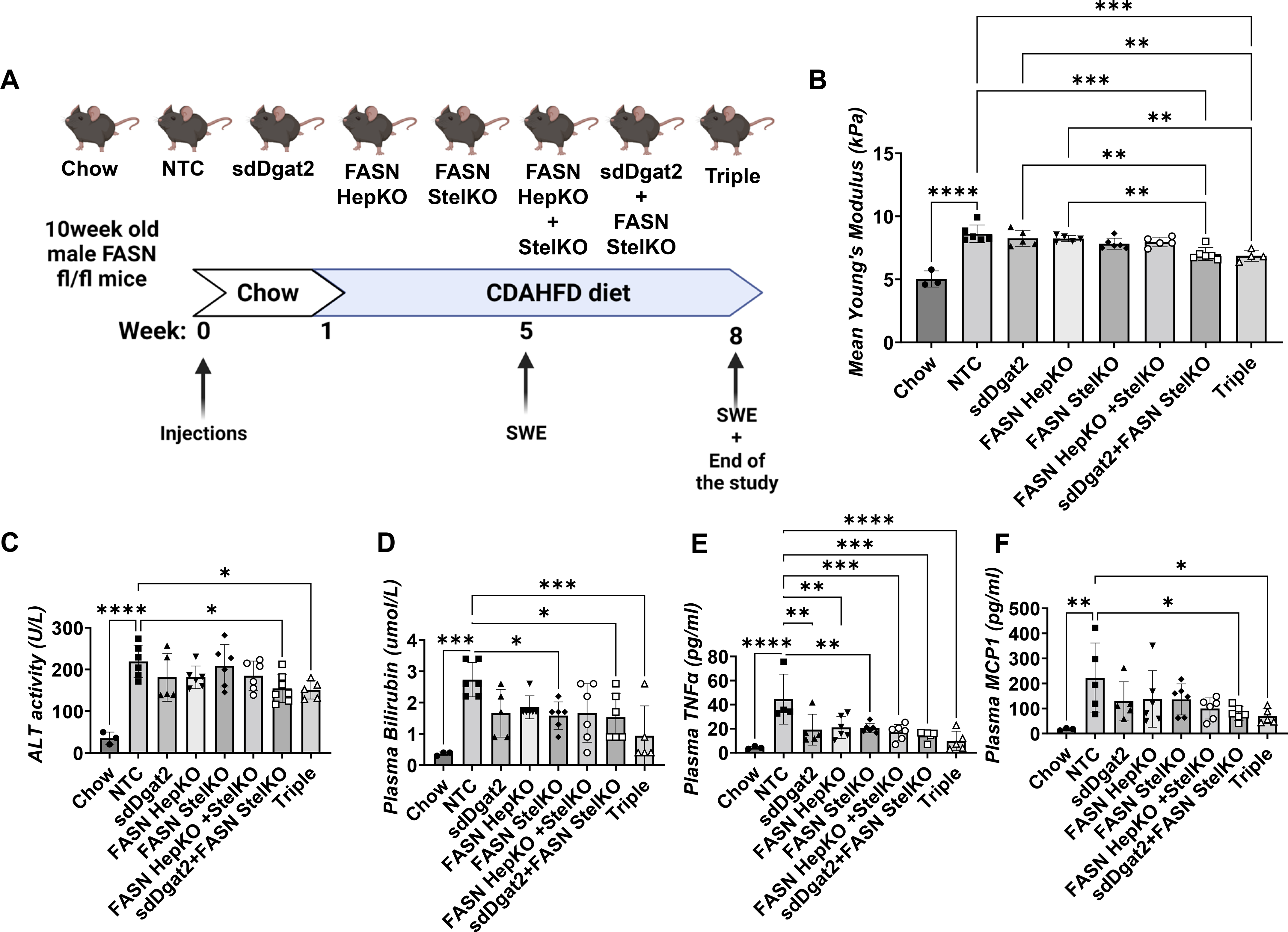
Stellate cell-specific deletion of FASN, in combination with DGAT2 silencing, ameliorates liver stiffness and plasma markers of NASH in CDAHFD-fed mice. **(A)** 10-week-old male Fasn fl/fl mice were injected with corresponding AAVs to obtain hepatocyte-specific KO (AAV8-TBG-Cre) and/or stellate cell-specific KO (AAV9-Lrat-Cre) via IV injection. Additionally, for the indicated groups, sdDgat2 (10mg/kg) was injected via a single subcutaneous injection. After one week of injections, mice were put on CDAHFD for 7 weeks. **(B)** Liver stiffness measurements by shear-wave elastography **(C)** Plasma ALT activity **(D)** Plasma total bilirubin **(E)** Plasma TNF alpha and **(F)** Plasma MCP1 protein levels. (ns: Not significant, *: p<0.05, **:p<0.005, ***:p<0.0005, ****:p<0.00005)

Shear wave elastography (SWE) enabled quantitative measurements of liver stiffness, a biomarker for estimating inflammation and fibrosis, without sacrificing the mice (Fig 3B). SWE was performed at week 5 as well as at the end of the study. At week 8, SWE measurements revealed that the combined depletions of FASN (in stellate cells or in stellate cells plus hepatocytes) plus DGAT2 in hepatocytes (sdDgat2+StelKO as well as sdDgat2+StelKO+HepKO, denoted Triple group) caused significantly less liver stiffness compared to NTC, sdDgat2, and FASN HepKO groups (Fig 3B). Furthermore, no stiffness difference was detected between the combined DGAT2 plus FASN depletion groups. Additionally, circulating indicators of liver damage and inflammation were measured, showing plasma ALT activity (Fig.3C) and bilirubin (Fig 3D) were significantly lower in mice that were depleted in both stellate cell FASN and DGAT2. Interestingly, all groups showed less plasma TNFα levels compared to the NTC group, suggesting depletions of either DGAT2 or FASN can attenuate inflammation by reducing lipotoxicity (Fig.3E). Interestingly, combined depletions of FASN plus DGAT2 significantly lowered plasma MCP1, while single depletions of FASN or DGAT2 did not (Fig.3F).

Analysis of livers from the mice described in Figure 2 showed *Dgat2* mRNA was strongly silenced by the sdDgat2 compound under all conditions in which it was administered (Figure 4A). *Fasn* mRNA levels were normal upon knockdown of *Dgat2* under the conditions of these experiments. Marked *Fasn* mRNA depletion was detected in livers of the FASN HepKO groups, but not in FASN StelKO groups, as stellate cells comprise only 5 to 8% of the whole liver. *Dgat2* mRNA was not affected in FASNKO groups in which sdDgat2 was not administered. In a separate study, we confirmed that the AAV9-LRAT-Cre construct efficiently and selectively depletes MCT1 in stellate cells when injected into MCT1^fl/fl^ mice^55^. Consistent with our previous results^24^, total liver triglycerides were significantly lowered in all sdDgat2 treated groups (Fig.4B). Immunohistochemical analysis of liver sections with an antibody against type 1 collagen revealed a striking prevention of fibrosis development in the Triple KO group (Fig.4C). Type 1 collagen quantified in liver sections from all mice in the NTC and Triple KO groups showed significantly lower collagen signal in the sdDgat2 plus FASN StelKO group and the Triple group compared to the NTC. Total hydroxyproline (OH-Pro) content was also significantly lower in these same two combinations DGAT2 plus FASN KO groups. Taken together, these results demonstrate a remarkable efficacy in preventing NASH development by combining FASNKO in liver stellate cells with DGAT2 depletion in hepatocytes in the CDHFD mouse model.

**Figure 4:**
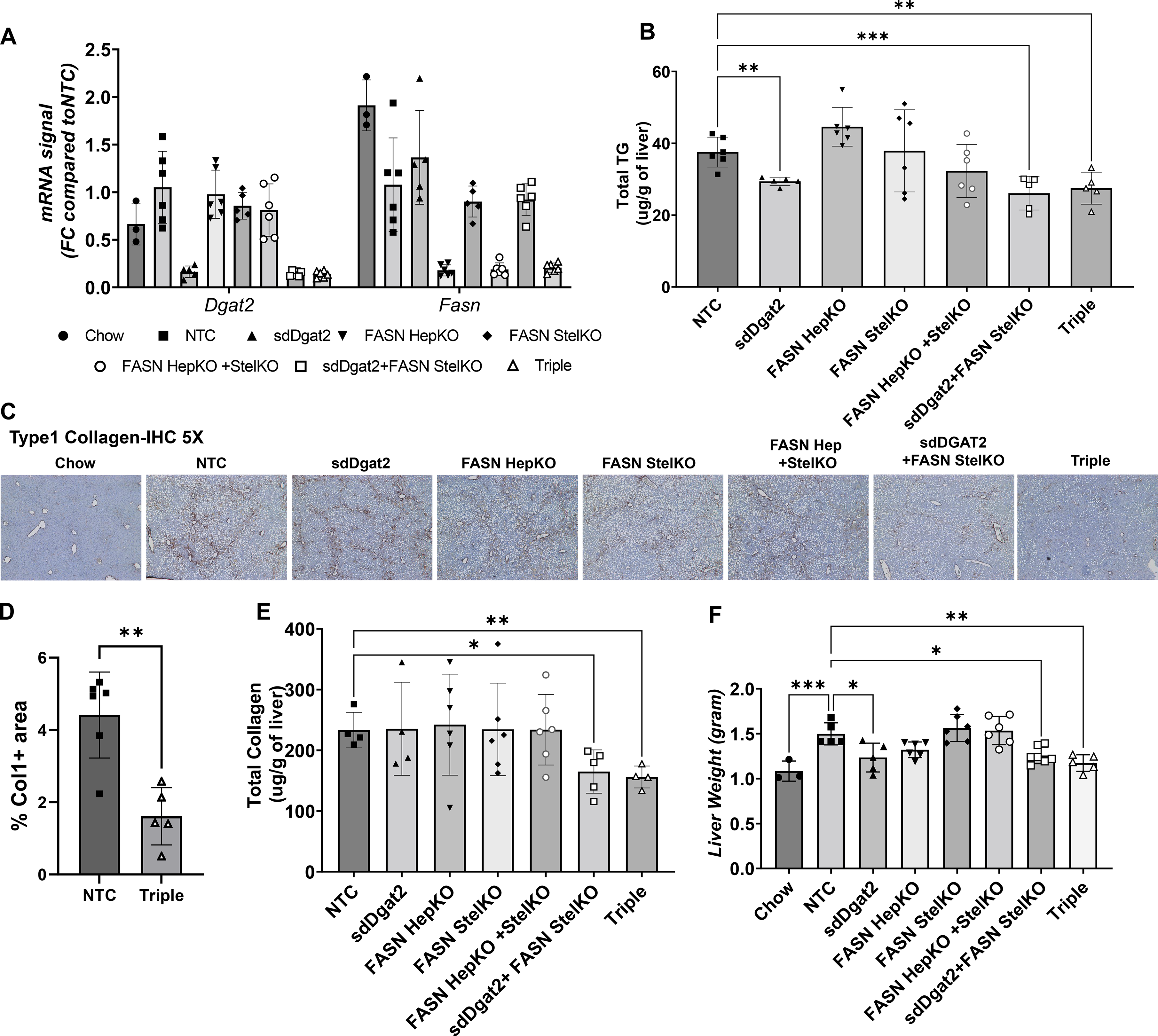
Stellate cell-specific deletion of FASN, in combination with DGAT2 silencing, prevents NASH in CDAHFD-fed mice. **(A)** mRNA levels of Dgat2 and Fasn **(B)** Total liver triglyceride measurement **(C)** Immunohistochemical analysis of type 1 collagen **(D)** Quantification of type 1 collagen positive area in IHC **(E)** Total collagen measurement via OH-Pro assay **(F)** Liver weights (ns: Not significant, *: p<0.05, **:p<0.005, ***:p<0.0005, ****:p<0.00005)

In summary, our studies reported here sought to address two major issues: 1) the mechanism by which DNL inhibition blocks fibrogenesis in stellate cells and 2) the contradictory results extrapolated from previous studies ^28, 37^ on FASN and DGAT2 single targeting strategies. Our findings reveal that the inhibition of autophagic flux is likely an underlying mechanism by which FASN depletion blocks fibrogenesis in stellate cells. Importantly, we also show that dual targeting hepatic steatosis in combination with the stellate fibrotic pathway is likely to be a stronger approach compared to monotherapy strategies. The fact that FASN depletion alone in stellate cells or in hepatocytes plus stellate cells does not alleviate progression of the fibrosis *in vivo* suggests that significant attenuation of hepatocyte steatosis is critical for FASN depletion to be maximally effective. Future studies are designed to optimize such dual targeting strategies for NASH therapeutics.

## Materials and Methods

### Human Stellate Cell Activation Studies

*In vitro* screening of LX2 human stellate cells was executed by plating 500,000 cells/well in 6-well plates overnight at 37°C, 5% CO2 in 0% FBS DMEM medium (11885-076, Gibco). Following a 4-hour pre-treatment with 1nM or 100 nM FASN inhibitor (TVB-3664-S8563, Selleck Chemicals), stellate cell activation was initiated by adding recombinant human Tgf-β (240-B-002, R & D Systems) for a 5ng/ml final concentration. After 48 hours, protein and mRNA were isolated from the cells and further analysis was performed.

### In vitro screening of chemically modified siRNAs

The initial in vitro screening was carried out by plating 20K cells/well in a 96 well plate. The cells were treated with the candidate compounds in final concentration of 1.5 µM. Cells were treated for 48 hours at 37 °C, 5% CO2 in 0% FBS DMEM medium. After 48hours, mRNA levels were quantified using the QuantiGene 2.0 assay kit (Affymetrix, QS0011. Briefly, cells were lysed in 300 µL diluted lysis mixture composed of one-part lysis mixture (Affymetrix, QG0503), 2 parts H2O, and 0.167 µg/µL proteinase K (ThermoScientific, EO0491) for 1 hour at 55 °C. Cell lysates were mixed thoroughly, and 20 µL/well of each lysate was added to the capture plate with 60 µL diluted lysis mixture without proteinase K. Probe sets for *FASN* and *B2M* (Affymetrix; 80246, 80003) were used according to the manufacturer’s recommended protocol. Datasets were normalized to B2M. The dose response assessments were done using serially diluted doses of the two candidate compounds (1.500, 0.750, 0.375, 0.188, 0.094, 0.047, 0.023, 0.012 µM), each condition in triplicate and the QuantiGene 2.0 Assay was performed.

### AAV Constructs

AAV8-TBG-Cre and Control AAV constructs were produced by UMass Chan Viral Core, as previously described ^56^. AAV9-Lrat-Cre was designed in house ^55^ and produced by Vector BioLabs.

### Animal studies

Animal experiments were performed in accordance with animal care ethics approval and guidelines of University of Massachusetts Medical School Institutional Animal Care and Use Committee (protocol number: A-1600-19). For *in vivo* studies, ten-week-old, male, FASN^fl/fl^ mice were each injected with either AAV8-TBG-Cre (hepatocyte specific KO), AAV9-Lrat-Cre (stellate cell specific KO), or a combination of the two constructs via intravenous injection. Each mouse was injected with 3 × 10^11^ GC/mL. Specifically, hepatocyte-specific KO mice received 1 × 10^11^ AAV8-TBG-Cre and 2 × 10^11^ Control AAV, stellate cell-specific KO received 2 × 10^11^ AAV9-LRAT-Cre and 1 × 10^11^ Control AAV, and the multiple-KO groups were given both 1 × 10^11^ AAV8-TBG-Cre and 2 × 10^11^ AAV9-LRAT-Cre. Following administration of the AAV constructs, the mice were placed on a Chow diet for one week before transitioning to HFD for the remaining 7 weeks. On weeks 5 and 8, liver stiffness was measured via shear-wave elastography.

### Oligonucleotide synthesis

Oligonucleotides were synthesized using modified (2ʹ-F, 2ʹ-O-Me) phosphoramidites with standard protecting groups. 5’-Vinyl Tetra phosphonate (pivaloyloxymethyl) 2’-O-Methyl Uridine 3’-CE phosphoramidite (VP) was used for the 5’-Vinyl-phosphonate coupling when needed. All amidites were purchased from (Chemgenes, Wilmington, MA). Phosphoramidite solid-phase synthesis was done on MerMade12 (Biosearch Technologies, Novato, CA) using modified protocols. GalNAc conjugated oligonucleotides were grown on a 500Å LCAA custom aminopropanediol-based trivalent GalNAc-CPG (Centernauchsnab, Minsk, Belarus). Phosphoramidites were prepared at 0.1 M in anhydrous acetonitrile (ACN), with added dry 15% dimethylformamide in the 2’-OMe-Uridine amidite. 5-(Benzylthio)-1H-tetrazole (BTT) was used as the activator at 0.25 M. Detritylations were performed using 3% trichloroacetic acid in dichloromethane. Capping reagents used were CAP A, 20% n-methylimidazole in ACN and CAP B, 20% acetic anhydride, 30% 2,6-lutidine in ACN (Synthesis reagents were purchased at AIC, Westborough, MA). Sulfurization was performed with 0.1 M solution of 3-[(dimethylaminomethylene)amino]-3H-1,2,4-dithiazole-5-thione (DDTT) in pyridine (Chemgenes, Wilmington, MA) for 3 min. Phosphoramidite coupling times were 4 min.

### Deprotection and purification of oligonucleotides

Conjugated oligonucleotides were cleaved and deprotected 28-30% ammonium hydroxide and 40% aq. methylamine (AMA) in a 1:1 ratio, for 2h at room temperature. The VP containing oligonucleotides were cleaved and deprotected as previously described ^57^. Briefly, CPG with VP-oligonucleotides was treated with a solution of 3% Diethylamine in 28-30% ammonium hydroxide at 35⁰C for 20h.

The solutions containing cleaved oligonucleotides were filtered to remove the CPG and dried under vacuum. The resulting pellets were re-suspended in 5% ACN in water. Purifications were performed on an Agilent 1290 Infinity II HPLC system. VP and GalNAc conjugated oligonucleotides were purified using a custom 20×150mm column packed with Source 15Q anion exchange resin (Cytiva, Marlborough, MA); run conditions: eluent A, 10 mM sodium acetate in 20% ACN in water; eluent B, 1 M sodium perchlorate in 20% ACN in water; linear gradient, 10 to 35% B 20 min at 40°C. Flow was 40mL/min and peaks were monitored at 260 nm. Fractions were analyzed by liquid chromatography mass spectrometry (LC–MS), pure fractions were dried under vacuum. Oligonucleotides were re-suspended in 5% ACN and desalted by size exclusion on a 50×250 mm custom column packed with Sephadex G-25 media (Cytiva, Marlborough, MA), and lyophilized.

### LC–MS analysis of oligonucleotides

The identity of oligonucleotides was verified by LC–MS analysis on an Agilent 6530 accurate mass Q-TOF using the following conditions: buffer A: 100 mM 1,1,1,3,3,3-hexafluoroisopropanol (HFIP) and 9 mM triethylamine (TEA) in LC–MS grade water; buffer B:100 mM HFIP and 9 mM TEA in LC–MS grade methanol; column, Agilent AdvanceBio oligonucleotides C18; linear gradient 0–30% B 8min (VP and GalNAc); temperature, 60°C; flow rate, 0.5 ml/min. LC peaks were monitored at 260nm. MS parameters: Source, electrospray ionization; ion polarity, negative mode; range, 100– 3,200 m/z; scan rate, 2 spectra/s; capillary voltage, 4,000; fragmentor, 180 V. Deprotection, purification and LC-MS reagents were purchased from Fisher Scientific, Sigma Aldrich and Oakwood Chemicals

### RNA isolation and RT–quantitative PCR

Frozen liver tissue punches (25–50 mg) were homogenized in TRIzol (15596018, Thermo Scientific) using the Qiagen TissueLyser II. Chloroform was added to the homogenate and centrifuged for 15 minutes at maximum speed. The clear upper layer was added to an equal volume of 100% isopropanol and incubated for 1 h at 4 °C. After 10 min of centrifugation at maximum speed, the supernatant was discarded, and 70 to 75% ethanol was added to wash the pellet. After 5 min of centrifugation at maximum speed, the supernatant was discarded, and the pellet was briefly dried in the hood prior to resuspension in double-distilled water. RNA concentration was measured via the Thermo Scientific NanoDrop2000 spectrophotometer. Complementary DNA was then synthesized from 1 μg of total RNA using iScript cDNA Synthesis Kit (1708890, Bio-Rad) and Bio-Rad T100 thermocycler. Quantitative RT–PCR was performed using iQ SybrGreen Supermix (4368708, Applied Biosystems) on the Bio-Rad CFX96 C1000 Touch Thermal Cycler and analyzed.

### Transmission electron microscopy

TEM analysis was carried out by the UMass Chan Medical School Cryo-EM Core. LX2 human stellate cells were plated 500k cells/well in 6-well plates overnight at 37°C, 5% CO2 in 0% FBS DMEM medium (11995065, Gibco). Following a 6-hour treatment with either Chol-FASN-2314, Chol-NTC or no treatment, stellate cell activation was initiated by adding recombinant human Tgf-β (240-B-002, R & D Systems) at a 5ng/ml final concentration. After 48 hours cells were fixed by first removing half the plate media and then adding an equal volume of 2.5% glutaraldehyde (v/v) in 1 M Na cacodylate buffer (pH 7.2) for 10 min before being transferred to 2.5% glutaraldehyde (v/v) in 1 M Na cacodylate buffer (pH 7.2) for 1 hour. Next, the cell plates were briefly rinsed (3 × 10 min) in 1 M Na cacodylate buffer (pH 7.2) and post-fixed for 1 hr in 1% osmium tetroxide (w/v) in dH2O. After washing three times with dH2O for 10 mins, cells were harvested with a soft plastic spatula, and pelleted by centrifugation. Samples were washed three times with dH2O for 10 min and dehydrated through a graded series of ethanol (10, 30, 50, 70, 85, 95% for 20 min each) to three changes of 100% ethanol. Samples were treated first with two changes of 100% Propylene Oxide and then with a 50%/50% propylene oxide/SPI-Pon 812 resin mixture overnight. The next morning the cell pellets were transferred through four changes of fresh SPI-pon 812-Araldite epoxy resin and finally embedded in tubes filled with the same resin and polymerized for 48 hr at 70 °C. The epoxy blocks were then trimmed, and ultrathin sections were cut on a Leica EM UC7 microtome using a diamond knife. The sections were collected and mounted on copper support grids and contrasted with lead citrate and uranyl acetate. The samples were examined on a Philips CM 10 using 100 Kv accelerating voltage. Images were captured using a Gatan TEM CCD camera.

### Immunoblotting

In the analysis of protein expression, cell lysates were homogenized in a RIPA buffer (25 mM Tris, pH 7.4 0.15 M NaCl 0.1% Tween 20) with 1:100 protease inhibitor (78442, Thermo Scientific). The lysates were denatured by boiling, separated on a 4 to 15% SDS/polyacrylamide gel electrophoresis gel (567-1094, Bio-Rad), and transferred to a nitrocellulose membrane (1704159, Bio-Rad). The membrane was blocked using TBS-T with 5% BSA for 1 hour at room temperature and incubated using p-SMAD3 (ab52903, Abcam), type I collagen (1310-30, Southern Biotech), and GAPDH (5174, Cell Signaling). The blot was further washed in TBS-T and incubated at room temperature with its corresponding secondary antibody for 30 minutes. Following another wash, the blot was incubated with ECL (32106, Thermo Scientific) and visualized with the ChemiDox XRS+ image-forming system.

### Plasma Measurements

Alanine transaminase levels were quantified using the Alanine Transaminase Colorimetric Activity Assay Kit® from Cayman Chemical. The instructions found in the user manual were followed utilizing plasma collected via retro orbital bleed prior to sacrifice. Absorbance was read by the Tecan safire2 microplate reader. Rest of the plasma measurements were done by UMass Chan Metabolic Core.

### Triglyceride Assay

Frozen liver punches (100-150 mg) were used to determine TG levels from tissue homogenates. Instructions found in the booklet of Cayman’s Triglyceride Colorimetric Assay® (10010303) were followed to quantify glycerol fluorometric measurements resulting from the enzymatic hydrolysis of triglycerides via lipase.

### Histological analysis

For the IHC, one lobe of the liver was fixed in 4% paraformaldehyde and embedded in paraffin. Sectioned slides were then stained type I collagen (Southern Biotech) at the UMass Medical School Morphology Core. Photos from the liver sections were taken with an Axiovert 35 Zeiss microscope (Zeiss) equipped with an Axiocam CCI camera at the indicated magnification.

### Collagen Assay

Total Collagen was measured from frozen liver punches (200-250 mg) using Quickzyme Total Collagen Assay® (QZBTOTCOL1). Instructions found in this Assay package were followed as it utilizes the presence of hydroxyproline to determine the amount of collagen present from tissue homogenates.

### Software & Statistics

All statistical analyses were performed using the GraphPad Prism 9 (GraphPad Software, Inc.). The data are presented as mean ± SEM. For analysis of the statistical significance between four or more groups, two-way ANOVA test was used. Ns is nonsignificant (p > 0.05), *:p < 0.05, **:p < 0.005, and ***:p < 0.0005.

## Acknowledgements

We wish to thank members of the Czech and Khvorova laboratories for helpful discussions, and Kerri Miller and Mary Beth Dziewietin for excellent assistance throughout the project. We thank the UMass Chan Medical School Morphology Core Facility for assistance in the histological preparations, stains, and analysis. We also thank the UMass Chan Medical School Cryo-EM Core Facility for their assistance in obtaining the TEM data. This work was supported by the National Institutes of Health grants DK103047 and DK116056 (to M.P.C.), R35GM131839 (to A.K.), S10OD020012 (to A.K.) and S10 OD025113-01(Cryo-EM Core Facility). We also gratefully acknowledge generous funding through the Isadore and Fannie Foxman Chair in Medical Science (to M.P.C.).

## Author contributions

B.Y. and M.P.C. designed the study, analyzed the data, and wrote the manuscript. B.Y., S.H., C.D, M.K., and K.M. performed most of the experiments and analyzed the data. D.E., B.M.B and S.O.J contributed to the synthesis of the oligonucleotides. A.K and D.E. provided guidance in oligonucleotide technology. J.K.K performed the plasma NASH marker analysis. K.R performed the transmission electron microscopy imaging.

## References

1. Younossi, Z.M., Blissett, D., Blissett, R., Henry, L., Stepanova, M., Younossi, Y., Racila, A., Hunt, S., and Beckerman, R. (2016). The economic and clinical burden of nonalcoholic fatty liver disease in the United States and Europe. Hepatology 64, 1577–1586. 10.1002/hep.28785.

2. Le, M.H., Yeo, Y.H., Zou, B., Barnet, S., Henry, L., Cheung, R., and Nguyen, M.H. (2022). Forecasted 2040 global prevalence of nonalcoholic fatty liver disease using hierarchical bayesian approach. Clin Mol Hepatol 28, 841–850. 10.3350/cmh.2022.0239.

3. Younossi, Z.M., Stepanova, M., Ong, J., Trimble, G., AlQahtani, S., Younossi, I., Ahmed, A., Racila, A., and Henry, L. (2021). Nonalcoholic Steatohepatitis Is the Most Rapidly Increasing Indication for Liver Transplantation in the United States. Clin Gastroenterol Hepatol 19, 580–589 e585. 10.1016/j.cgh.2020.05.064.

4. Alexander, M., Loomis, A.K., van der Lei, J., Duarte-Salles, T., Prieto-Alhambra, D., Ansell, D., Pasqua, A., Lapi, F., Rijnbeek, P., Mosseveld, M., et al. (2019). Risks and clinical predictors of cirrhosis and hepatocellular carcinoma diagnoses in adults with diagnosed NAFLD: real-world study of 18 million patients in four European cohorts. BMC Med 17, 95. 10.1186/s12916-019-1321-x.

5. Diehl, A.M., and Day, C. (2017). Cause, Pathogenesis, and Treatment of Nonalcoholic Steatohepatitis. N Engl J Med 377, 2063–2072. 10.1056/NEJMra1503519.

6. Esler, W.P., and Bence, K.K. (2019). Metabolic Targets in Nonalcoholic Fatty Liver Disease. Cell Mol Gastroenterol Hepatol 8, 247–267. 10.1016/j.jcmgh.2019.04.007.

7. Lomonaco, R., Ortiz-Lopez, C., Orsak, B., Webb, A., Hardies, J., Darland, C., Finch, J., Gastaldelli, A., Harrison, S., Tio, F., and Cusi, K. (2012). Effect of adipose tissue insulin resistance on metabolic parameters and liver histology in obese patients with nonalcoholic fatty liver disease. Hepatology 55, 1389–1397. 10.1002/hep.25539.

8. Noureddin, M., and Sanyal, A.J. (2018). Pathogenesis of NASH: The Impact of Multiple Pathways. Curr Hepatol Rep 17, 350–360. 10.1007/s11901-018-0425-7.

9. Younossi, Z.M., Koenig, A.B., Abdelatif, D., Fazel, Y., Henry, L., and Wymer, M. (2016). Global epidemiology of nonalcoholic fatty liver disease-Meta-analytic assessment of prevalence, incidence, and outcomes. Hepatology 64, 73–84. 10.1002/hep.28431.

10. Friedman, S.L., Neuschwander-Tetri, B.A., Rinella, M., and Sanyal, A.J. (2018). Mechanisms of NAFLD development and therapeutic strategies. Nat Med 24, 908–922. 10.1038/s41591-018-0104-9.

11. Ganz, M., and Szabo, G. (2013). Immune and inflammatory pathways in NASH. Hepatol Int 7 Suppl 2, 771–781. 10.1007/s12072-013-9468-6.

12. Llovet, J.M., Willoughby, C.E., Singal, A.G., Greten, T.F., Heikenwalder, M., El-Serag, H.B., Finn, R.S., and Friedman, S.L. (2023). Nonalcoholic steatohepatitis-related hepatocellular carcinoma: pathogenesis and treatment. Nat Rev Gastroenterol Hepatol. 10.1038/s41575-023-00754-7.

13. Muthiah, M.D., and Sanyal, A.J. (2020). Current management of non-alcoholic steatohepatitis. Liver Int 40 Suppl 1, 89–95. 10.1111/liv.14355.

14. Tilg, H., Adolph, T.E., and Moschen, A.R. (2021). Multiple Parallel Hits Hypothesis in Nonalcoholic Fatty Liver Disease: Revisited After a Decade. Hepatology 73, 833–842. 10.1002/hep.31518.

15. Wong, V.W., and Singal, A.K. (2019). Emerging medical therapies for non-alcoholic fatty liver disease and for alcoholic hepatitis. Transl Gastroenterol Hepatol 4, 53. 10.21037/tgh.2019.06.06.

16. Eslam, M., and George, J. (2020). Genetic contributions to NAFLD: leveraging shared genetics to uncover systems biology. Nat Rev Gastroenterol Hepatol 17, 40–52. 10.1038/s41575-019-0212-0.

17. Koo, B.K., Joo, S.K., Kim, D., Bae, J.M., Park, J.H., Kim, J.H., and Kim, W. (2018). Additive effects of PNPLA3 and TM6SF2 on the histological severity of non-alcoholic fatty liver disease. J Gastroenterol Hepatol 33, 1277–1285. 10.1111/jgh.14056.

18. Kozlitina, J., Smagris, E., Stender, S., Nordestgaard, B.G., Zhou, H.H., Tybjaerg-Hansen, A., Vogt, T.F., Hobbs, H.H., and Cohen, J.C. (2014). Exome-wide association study identifies a TM6SF2 variant that confers susceptibility to nonalcoholic fatty liver disease. Nat Genet 46, 352–356. 10.1038/ng.2901.

19. Kozlitina, J., Stender, S., Hobbs, H.H., and Cohen, J.C. (2018). HSD17B13 and Chronic Liver Disease in Blacks and Hispanics. N Engl J Med 379, 1876–1877. 10.1056/NEJMc1804027.

20. Mancina, R.M., Dongiovanni, P., Petta, S., Pingitore, P., Meroni, M., Rametta, R., Boren, J., Montalcini, T., Pujia, A., Wiklund, O., et al. (2016). The MBOAT7-TMC4 Variant rs641738 Increases Risk of Nonalcoholic Fatty Liver Disease in Individuals of European Descent. Gastroenterology 150, 1219–1230 e1216. 10.1053/j.gastro.2016.01.032.

21. Romeo, S., Sanyal, A., and Valenti, L. (2020). Leveraging Human Genetics to Identify Potential New Treatments for Fatty Liver Disease. Cell Metab 31, 35–45. 10.1016/j.cmet.2019.12.002.

22. Speliotes, E.K., Yerges-Armstrong, L.M., Wu, J., Hernaez, R., Kim, L.J., Palmer, C.D., Gudnason, V., Eiriksdottir, G., Garcia, M.E., Launer, L.J., et al. (2011). Genome-wide association analysis identifies variants associated with nonalcoholic fatty liver disease that have distinct effects on metabolic traits. PLoS Genet 7, e1001324. 10.1371/journal.pgen.1001324.

23. Valenti, L., Al-Serri, A., Daly, A.K., Galmozzi, E., Rametta, R., Dongiovanni, P., Nobili, V., Mozzi, E., Roviaro, G., Vanni, E., et al. (2010). Homozygosity for the patatin-like phospholipase-3/adiponutrin I148M polymorphism influences liver fibrosis in patients with nonalcoholic fatty liver disease. Hepatology 51, 1209–1217. 10.1002/hep.23622.

24. Yenilmez, B., Wetoska, N., Kelly, M., Echeverria, D., Min, K., Lifshitz, L., Alterman, J.F., Hassler, M.R., Hildebrand, S., DiMarzio, C., et al. (2022). An RNAi therapeutic targeting hepatic DGAT2 in a genetically obese mouse model of nonalcoholic steatohepatitis. Mol Ther 30, 1329–1342. 10.1016/j.ymthe.2021.11.007.

25. Amin, N.B., Carvajal-Gonzalez, S., Purkal, J., Zhu, T., Crowley, C., Perez, S., Chidsey, K., Kim, A.M., and Goodwin, B. (2019). Targeting diacylglycerol acyltransferase 2 for the treatment of nonalcoholic steatohepatitis. Sci Transl Med 11. 10.1126/scitranslmed.aav9701.

26. Calle, R.A., Amin, N.B., Carvajal-Gonzalez, S., Ross, T.T., Bergman, A., Aggarwal, S., Crowley, C., Rinaldi, A., Mancuso, J., Aggarwal, N., et al. (2021). ACC inhibitor alone or co-administered with a DGAT2 inhibitor in patients with non-alcoholic fatty liver disease: two parallel, placebo-controlled, randomized phase 2a trials. Nat Med 27, 1836–1848. 10.1038/s41591-021-01489-1.

27. Choi, C.S., Savage, D.B., Kulkarni, A., Yu, X.X., Liu, Z.X., Morino, K., Kim, S., Distefano, A., Samuel, V.T., Neschen, S., et al. (2007). Suppression of diacylglycerol acyltransferase-2 (DGAT2), but not DGAT1, with antisense oligonucleotides reverses diet-induced hepatic steatosis and insulin resistance. J Biol Chem 282, 22678–22688. 10.1074/jbc.M704213200.

28. Loomba, R., Morgan, E., Watts, L., Xia, S., Hannan, L.A., Geary, R.S., Baker, B.F., and Bhanot, S. (2020). Novel antisense inhibition of diacylglycerol O-acyltransferase 2 for treatment of non-alcoholic fatty liver disease: a multicentre, double-blind, randomised, placebo-controlled phase 2 trial. Lancet Gastroenterol Hepatol 5, 829–838. 10.1016/S2468-1253(20)30186-2.

29. Yamaguchi, K., Yang, L., McCall, S., Huang, J., Yu, X.X., Pandey, S.K., Bhanot, S., Monia, B.P., Li, Y.X., and Diehl, A.M. (2007). Inhibiting triglyceride synthesis improves hepatic steatosis but exacerbates liver damage and fibrosis in obese mice with nonalcoholic steatohepatitis. Hepatology 45, 1366–1374. 10.1002/hep.21655.

30. Yu, X.X., Murray, S.F., Pandey, S.K., Booten, S.L., Bao, D., Song, X.Z., Kelly, S., Chen, S., McKay, R., Monia, B.P., and Bhanot, S. (2005). Antisense oligonucleotide reduction of DGAT2 expression improves hepatic steatosis and hyperlipidemia in obese mice. Hepatology 42, 362–371. 10.1002/hep.20783.

31. Yen, C.L., Stone, S.J., Koliwad, S., Harris, C., and Farese, R.V., Jr. (2008). Thematic review series: glycerolipids. DGAT enzymes and triacylglycerol biosynthesis. J Lipid Res 49, 2283–2301. 10.1194/jlr.R800018-JLR200.

32. Zammit, V.A. (2013). Hepatic triacylglycerol synthesis and secretion: DGAT2 as the link between glycaemia and triglyceridaemia. Biochem J 451, 1–12. 10.1042/BJ20121689.

33. Choi, S., and Diehl, A.M. (2005). Role of inflammation in nonalcoholic steatohepatitis. Curr Opin Gastroenterol 21, 702–707. 10.1097/01.mog.0000182863.96421.47.

34. Loomba, R., Noureddin, M., Kowdley, K.V., Kohli, A., Sheikh, A., Neff, G., Bhandari, B.R., Gunn, N., Caldwell, S.H., Goodman, Z., et al. (2021). Combination Therapies Including Cilofexor and Firsocostat for Bridging Fibrosis and Cirrhosis Attributable to NASH. Hepatology 73, 625–643. 10.1002/hep.31622.

35. Bates, J., Vijayakumar, A., Ghoshal, S., Marchand, B., Yi, S., Kornyeyev, D., Zagorska, A., Hollenback, D., Walker, K., Liu, K., et al. (2020). Acetyl-CoA carboxylase inhibition disrupts metabolic reprogramming during hepatic stellate cell activation. J Hepatol 73, 896–905. 10.1016/j.jhep.2020.04.037.

36. O’Farrell, M., Duke, G., Crowley, R., Buckley, D., Martins, E.B., Bhattacharya, D., Friedman, S.L., and Kemble, G. (2022). FASN inhibition targets multiple drivers of NASH by reducing steatosis, inflammation and fibrosis in preclinical models. Sci Rep 12, 15661. 10.1038/s41598-022-19459-z.

37. Loomba, R., Mohseni, R., Lucas, K.J., Gutierrez, J.A., Perry, R.G., Trotter, J.F., Rahimi, R.S., Harrison, S.A., Ajmera, V., Wayne, J.D., et al. (2021). TVB-2640 (FASN Inhibitor) for the Treatment of Nonalcoholic Steatohepatitis: FASCINATE-1, a Randomized, Placebo-Controlled Phase 2a Trial. Gastroenterology 161, 1475–1486. 10.1053/j.gastro.2021.07.025.

38. Syed-Abdul, M.M., Parks, E.J., Gaballah, A.H., Bingham, K., Hammoud, G.M., Kemble, G., Buckley, D., McCulloch, W., and Manrique-Acevedo, C. (2020). Fatty Acid Synthase Inhibitor TVB-2640 Reduces Hepatic de Novo Lipogenesis in Males With Metabolic Abnormalities. Hepatology 72, 103–118. 10.1002/hep.31000.

39. Rowland, L.A., Guilherme, A., Henriques, F., DiMarzio, C., Munroe, S., Wetoska, N., Kelly, M., Reddig, K., Hendricks, G., Pan, M., et al. (2023). De novo lipogenesis fuels adipocyte autophagosome and lysosome membrane dynamics. Nat Commun 14, 1362. 10.1038/s41467-023-37016-8.

40. Hernandez-Gea, V., Ghiassi-Nejad, Z., Rozenfeld, R., Gordon, R., Fiel, M.I., Yue, Z., Czaja, M.J., and Friedman, S.L. (2012). Autophagy releases lipid that promotes fibrogenesis by activated hepatic stellate cells in mice and in human tissues. Gastroenterology 142, 938–946. 10.1053/j.gastro.2011.12.044.

41. Lucantoni, F., Martinez-Cerezuela, A., Gruevska, A., Moragrega, A.B., Victor, V.M., Esplugues, J.V., Blas-Garcia, A., and Apostolova, N. (2021). Understanding the implication of autophagy in the activation of hepatic stellate cells in liver fibrosis: are we there yet? J Pathol 254, 216–228. 10.1002/path.5678.

42. Thoen, L.F., Guimaraes, E.L., Dolle, L., Mannaerts, I., Najimi, M., Sokal, E., and van Grunsven, L.A. (2011). A role for autophagy during hepatic stellate cell activation. J Hepatol 55, 1353–1360. 10.1016/j.jhep.2011.07.010.

43. Trivedi, P., Wang, S., and Friedman, S.L. (2021). The Power of Plasticity-Metabolic Regulation of Hepatic Stellate Cells. Cell Metab 33, 242–257. 10.1016/j.cmet.2020.10.026.

44. Afify, H., Ghoneum, A., Almousa, S., Abdulfattah, A.Y., Warren, B., Langsten, K., Gonzalez, D., Casals, R., Bharadwaj, M., Kridel, S., and Said, N. (2021). Metabolomic credentialing of murine carcinogen-induced urothelial cancer. Sci Rep 11, 22085. 10.1038/s41598-021-99746-3.

45. Chu, J., Xing, C., Du, Y., Duan, T., Liu, S., Zhang, P., Cheng, C., Henley, J., Liu, X., Qian, C., et al. (2021). Pharmacological inhibition of fatty acid synthesis blocks SARS-CoV-2 replication. Nat Metab 3, 1466–1475. 10.1038/s42255-021-00479-4.

46. Friedman, S.L. (2008). Hepatic stellate cells: protean, multifunctional, and enigmatic cells of the liver. Physiol Rev 88, 125–172. 10.1152/physrev.00013.2007.

47. Fondevila, M.F., Fernandez, U., Heras, V., Parracho, T., Gonzalez-Rellan, M.J., Novoa, E., Porteiro, B., Alonso, C., Mayo, R., da Silva Lima, N., et al. (2022). Inhibition of carnitine palmitoyltransferase 1A in hepatic stellate cells protects against fibrosis. J Hepatol 77, 15–28. 10.1016/j.jhep.2022.02.003.

48. Brown, K.E., Broadhurst, K.A., Mathahs, M.M., Brunt, E.M., and Schmidt, W.N. (2005). Expression of HSP47, a collagen-specific chaperone, in normal and diseased human liver. Lab Invest 85, 789–797. 10.1038/labinvest.3700271.

49. Prestigiacomo, V., and Suter-Dick, L. (2018). Nrf2 protects stellate cells from Smad-dependent cell activation. PLoS One 13, e0201044. 10.1371/journal.pone.0201044.

50. Nisimoto, Y., Diebold, B.A., Cosentino-Gomes, D., and Lambeth, J.D. (2014). Nox4: a hydrogen peroxide-generating oxygen sensor. Biochemistry 53, 5111–5120. 10.1021/bi500331y.

51. Alterman, J.F., Hall, L.M., Coles, A.H., Hassler, M.R., Didiot, M.C., Chase, K., Abraham, J., Sottosanti, E., Johnson, E., Sapp, E., et al. (2015). Hydrophobically Modified siRNAs Silence Huntingtin mRNA in Primary Neurons and Mouse Brain. Mol Ther Nucleic Acids 4, e266. 10.1038/mtna.2015.38.

52. Haraszti, R.A., Roux, L., Coles, A.H., Turanov, A.A., Alterman, J.F., Echeverria, D., Godinho, B., Aronin, N., and Khvorova, A. (2017). 5’-Vinylphosphonate improves tissue accumulation and efficacy of conjugated siRNAs in vivo. Nucleic Acids Res 45, 7581–7592. 10.1093/nar/gkx507.

53. Hassler, M.R., Turanov, A.A., Alterman, J.F., Haraszti, R.A., Coles, A.H., Osborn, M.F., Echeverria, D., Nikan, M., Salomon, W.E., Roux, L., et al. (2018). Comparison of partially and fully chemically-modified siRNA in conjugate-mediated delivery in vivo. Nucleic Acids Res 46, 2185–2196. 10.1093/nar/gky037.

54. Loos, B., du Toit, A., and Hofmeyr, J.H. (2014). Defining and measuring autophagosome flux-concept and reality. Autophagy 10, 2087–2096. 10.4161/15548627.2014.973338.

55. Min, K., Yenilmez, B., Kelly, M., Echeverria, D., Elleby, M., Lifshitz, L.M., Raymond, N., Tsagkaraki, E., Harney, S.M., DiMarzio, C., et al. (2023). Lactate transporter MCT1 in hepatic stellate cells promotes fibrotic collagen expression in nonalcoholic steatohepatitis. bioRxiv. 10.1101/2023.05.03.539244.

56. Sena-Esteves, M., and Gao, G. (2020). Introducing Genes into Mammalian Cells: Viral Vectors. Cold Spring Harb Protoc 2020, 095513. 10.1101/pdb.top095513.

57. O’Shea, J., Theile, C.S., Das, R., Babu, I.R., Charisse, K., Manoharan, M., Maier, M.A., and Zlatev, I. (2018). An efficient deprotection method for 5′-[O,O-bis(pivaloyloxymethyl)]-(E)-vinylphosphonate containing oligonucleotides. Tetrahedron 74, 6182–6186. https://doi.org/10.1016/j.tet.2018.09.008.

